# Association between the 4p16 genomic locus and different types of congenital heart disease: results from adult survivors in the UK Biobank

**DOI:** 10.1101/760371

**Authors:** Aldo Córdova-Palomera, James R. Priest

## Abstract

Congenital heart disease is the most common birth defect in newborns and the leading cause of death in infancy, affecting nearly 1% of live births. A locus in chromosome 4p16, adjacent to *MSX1* and *STX18*, has been associated with atrial septal defects (ASD) in multiple European and Chinese cohorts. Here, genotyping data from the UK Biobank was used to test for associations between this locus and congenital heart disease in adult survivors of left ventricular outflow tract obstruction (*n*=164) and ASD (*n*=223), with a control sample of 332,788 individuals, and a meta-analysis of the new and existing ASD data was performed.

The results show an association between the previously reported markers at 4p16 and risk for either ASD or left ventricular outflow tract obstruction, with effect sizes similar to the published data (OR between 1.27-1.45; all *p*<0.05). Differences in allele frequencies remained constant through the studied age range (40-70 years), indicating that the variants themselves do not drive lethal genetic defects. Meta-analysis shows an OR of 1.35 (95% CI: 1.25-1.46; *p*<10^−4^) for the association with ASD.

The findings show that the genetic associations with ASD can be generalized to adult survivors of both ASD and left ventricular lesions. Although the 4p16 associations are statistically compelling, the mentioned alleles confer only a small risk for disease and their frequencies in this adult sample are the same as in children, likely limiting their clinical significance. Further epidemiological and functional studies may elicit factors triggering disease in interaction with the risk alleles.

## INTRODUCTION

Affecting nearly 1% of live births, congenital heart disease (CHD) is the most common congenital anomaly in newborns and the principal cause of death in infancy ^1^. Expected long-term survival rates have considerably raised with advances in diagnostic and surgical techniques ^2,3^; survivors require lifelong treatment, posing a growing resource challenge for healthcare systems ^2^. Quantitative genetic research supports a strong genetic contribution to CHD ^4^, and epidemiological findings on survivors have evidenced multiple cardiac and non-cardiac comorbidities that pose novel treatment challenges.

Established molecular genetic contributions to CHD include aneuploidies, copy number variants, point and de novo mutations ^4,5^. Building on the current knowledge on CHD genetics, recent large-scale genome-wide association studies (GWAS) have identified some genetic variants increasing the risk for CHD-related phenotypes. In the current GWAS literature, the single most robust association between common genetic variation and CHD risk is a locus in chromosome 4p16, which was initially shown to confer risk for atrial septal defect (ASD) ^6^. This finding was originally reported by Cordell et al. ^6^ in two European-ancestry samples: a discovery cohort of 340 individuals with ostium secundum ASD and 5,159 controls, and a further 417 secundum ASD cases with 2,520 controls. Independent studies in Chinese cohorts have provided additional support to that finding: Zhao et al. ^7^ replicated the results in Han Chinese subjects (two cohorts, totaling 701 ASD cases and 3,208 controls); next, a similar outcome was observed in a sample from Southwest China (190 ASD cases and 225 controls) ^8^; and more recently, Pei et al. ^9^ reported analogous findings in a Fujian Chinese population (354 non-syndromic ASD cases and 557 controls).

The six samples mentioned above (two European- and four Chinese-ancestry) thus show consistent evidence of an association between the 4p16 locus and the ASD subtype of CHD -but no clear link to other CHD types–. These observations were based on datasets from either pediatric patients or young adults (median ages up to ~20 years). In this study, the 4p16 susceptibility locus is tested in relation to CHD in older adult survivors: left ventricular outflow tract obstruction (LVOTO) and ASD in a harmonized set of European-ancestry individuals, via the UK Biobank (164 LVOTO cases, 223 ASD cases and 332,788 non-CHD controls; median age of 58 years).

## METHODS

The dataset included in the adult CHD analyses was extracted from the UK Biobank, a longitudinal cohort study of over 500,000 individuals aged 40-69 years. Between 2006 and 2010, participants were enrolled in study centers across England, Scotland and Wales, and extensive baseline data on health behavior, medical history and physical measurements were collected through clinical examination and questionnaires. Similarly, blood, urine and saliva samples were collected from the participants. Ethical approval for the UK Biobank was granted by the North West Multi-Centre Research Ethics Committee and the National Health Service (NHS) National Research Ethics Service (ref: 11/NW/0382). All participants provided written informed consent to participate in the UK Biobank study, and all experiments were performed in agreement with relevant regulations and guidelines. Additional information, including details about the study protocol, is available online (https://biobank.ctsu.ox.ac.uk/).

CHD definitions and exclusion criteria were elaborated by leveraging information from medical histories, International Classification of Disease 9 and 10 (ICD-9 and 10) and Office of Population and Censuses Surveys (OPCS-4) diagnosis codes, in addition to information on age at diagnosis and age at surgery. The phenotype classification procedure consisted of four main steps: identification of all possible cases based on diagnostic and procedural codes; exclusion of unconfirmed cases for those who met criteria for non-congenital etiologies of heart disease (e.g., rheumatic); confirmation of case status for those whose age at diagnose suggested high likelihood of congenital etiology, or exclusion otherwise; and exclusion of control group for participants with CHD-related conditions due to lack of confidence to ascertain their status. More details about phenotype definitions are included in Supplementary Methods and Supplementary Figure S1.

Demographic data on sex and age distributions across diagnoses is summarized in Supplementary Table S1, organized as “controls” (subjects with no evidence of CHD), “LVOTO” (including both LVOTO and bicuspid aortic valve, BAV), “any ASD” (comprising participants with either a pure ASD diagnose, or an ASD and another CHD condition), or “pure ASD” (a subset of “any ASD”: people with evidence of ASD, but no other CHD). All disease categories (“LVOTO”, “any ASD” or “pure ASD”) had a higher number of male than female individuals, as opposed to the control subsample (statistically significant differences only for LVOTO versus controls). Age distributions were balanced across categories, ranging between 39 and 72 years of age in controls, and between 40 and 70 years in all CHD groups. As shown in Supplementary Table S2, all markers had minor allele frequency (MAF) between 0.238 and 0.239, in agreement with reference data, and the genotype counts in healthy participants did not depart from Hardy-Weinberg equilibrium. Within the sample of European-ancestry participants included in the analyses, pairwise estimates of linkage disequilibrium across all four markers were high (*R*^2^ between 0.969 and 0.994, as estimated by PLINK 1.9’s *--r2* command). The previous observation is in line with the presence of variants within a 33,767-base interval, previous and current association results on any of the four markers may thus represent a single signal. To avoid discussion of redundant analyses, the main marker reported by Cordell et al. ^6^ (rs870142) was used as tag SNP in the meta-analysis moving forward.

## RESULTS AND DISCUSSION

Power calculations using CaTS ^10^ indicated that, under an additive model with a prevalence of 1%, alpha of 0.05 and MAF of 23.8%, power to detect an effect size of 1.46 (similar magnitude to Cordell et al.’s ^6^ pooled European-ancestry data) was 88.6% for the UK Biobank’s LVOTO group, and 77.5% for the ASD phenotype. Table 1 shows statistics from Firth’s regression tests for the included markers. As displayed, the odds ratio (OR) for LVOTO conferred by the 4p16 genotypes was between 1.27 and 1.3 across all four markers. In the independent set of 223 ASD cases with and without other CHD comorbidities, effect sizes for the risk alleles were higher (ORs around 1.45), similar to the subset of ASD CHD with no comorbidities (*n*=167 cases from the larger 223-individual set, ORs between 1.47 and 1.5). Potential survival bias effects were tested by means of logistic regression with the addition of a genotype × age interaction term. As indicated in Figure 1 and Supplementary Figure S2, results of this analysis do not show evidence of age-modulating effects on the genotype-CHD associations, even though the distribution of ASD cases carrying the risk alleles (CT or TT genotypes) showed a consistent shift towards lower age (probably indicative of slightly younger ages at death).

**Table 1.** Association test results from additive models

|  |  |  |  |  | LVOTO |  |  |  | Any ASD<sup>1</sup> |  |  |  | Pure ASD<sup>1</sup> |  |  |  |
| --- | --- | --- | --- | --- | --- | --- | --- | --- | --- | --- | --- | --- | --- | --- | --- | --- |
| Chr. | Position | rsID | A1 | A2 | Beta | SE | *p* | FDR *p* | Beta | SE | *p* | FDR *p* | Beta | SE | *p* | FDR *p* |
| 4 | 4614280 | rs6824295 | T | C | 0.236 | 0.12 | 0.049 | 0.049 | 0.374 | 0.101 | 2×10<sup>-4</sup> | 2.1×10<sup>-4</sup> | 0.403 | 0.115 | 4.4×10<sup>-4</sup> | 6.6×10<sup>-4</sup> |
| 4 | 4628054 | rs4689904 | A | G | 0.25 | 0.119 | 0.036 | 0.048 | 0.372 | 0.1 | 2.1×10<sup>-4</sup> | 2.1×10<sup>-4</sup> | 0.4 | 0.115 | 4.9×10<sup>-4</sup> | 6.6×10<sup>-4</sup> |
| 4 | 4635276 | rs16835979 | A | C | 0.263 | 0.119 | 0.027 | 0.048 | 0.373 | 0.1 | 2×10<sup>-4</sup> | 2.1×10<sup>-4</sup> | 0.388 | 0.115 | 7.3×10<sup>-4</sup> | 7.4×10<sup>-4</sup> |
| 4 | 4648047 | rs870142 | T | C | 0.255 | 0.119 | 0.033 | 0.048 | 0.375 | 0.101 | 1.9×10<sup>-4</sup> | 2.1×10<sup>-4</sup> | 0.402 | 0.115 | 4.7×10<sup>-4</sup> | 6.6×10<sup>-4</sup> |

Notes: <sup>1</sup>, the “pure ASD” category includes individuals with ASD only, and is a subset of the participants in “any ASD”. Abbreviations: Chr., chromosome; rsID, SNP identifier; FDR *p*, false discovery rate-adjusted *p*-value.

**Figure 1.**
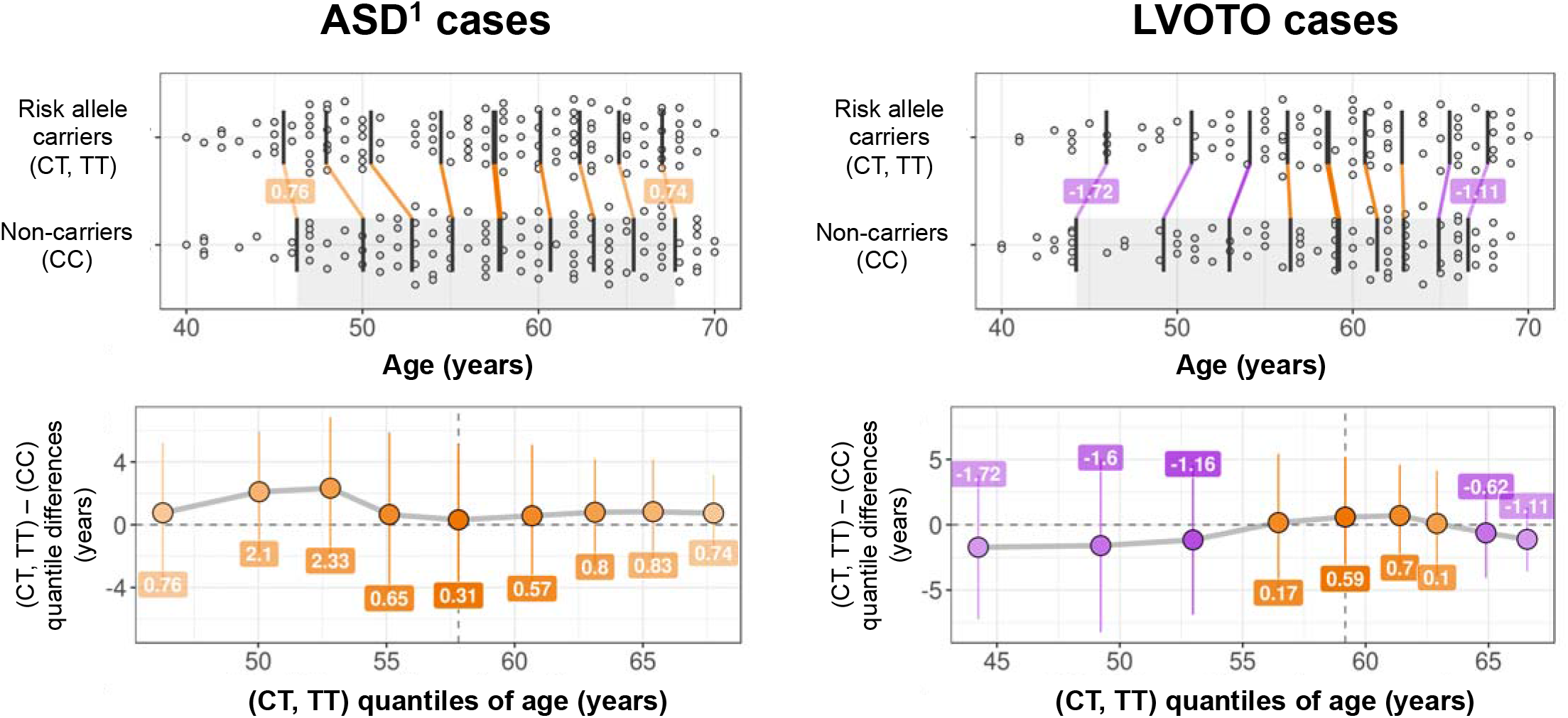
Shift function plots for age and rs870142 genotype for ASD and LVOTO cases. Top: Jittered marginal distribution scatterplots with overlaid shift function indicators using deciles, for “any ASD” (left) and LVOTO (right). 95% confidence intervals were computed using a percentile bootstrap estimation of the standard error of the difference between quantiles on 1000 bootstrap samples. Bottom: Linked deciles from shift functions. Note: ^1^, plotted ASD cases on the leftmost column correspond to the “any ASD” group, which includes ASD cases with or without other CHDs. Plots grouped using three genotypes (CC, CT and TT) are included in Supplementary Figure S2.

A meta-analysis of ASD was conducted using existing data and the new UK Biobank ASD results. The outcome of a random-effects meta-analysis model of the seven available datasets showed an overall OR of 1.35 (95% confidence interval: 1.25-1.46; *p*<10^−4^) (Figure 2). There was no statistical evidence of between-study heterogeneity (*I*^2^: 0%; *H*^2^: 1; *τ*^2^: 0; Q: 4.9; *p*_Q_=0.554). Publication bias was not observed in either a standard funnel plot (Figure 2, left) or a regression test for asymmetry (*z*=1.5, *p*=0.144). However, the contour-enhanced funnel plot (Figure 2, right) shows the absence of small effect sizes with significance levels below 95%, which might be indicative of biases other than publication bias.

**Figure 2.**
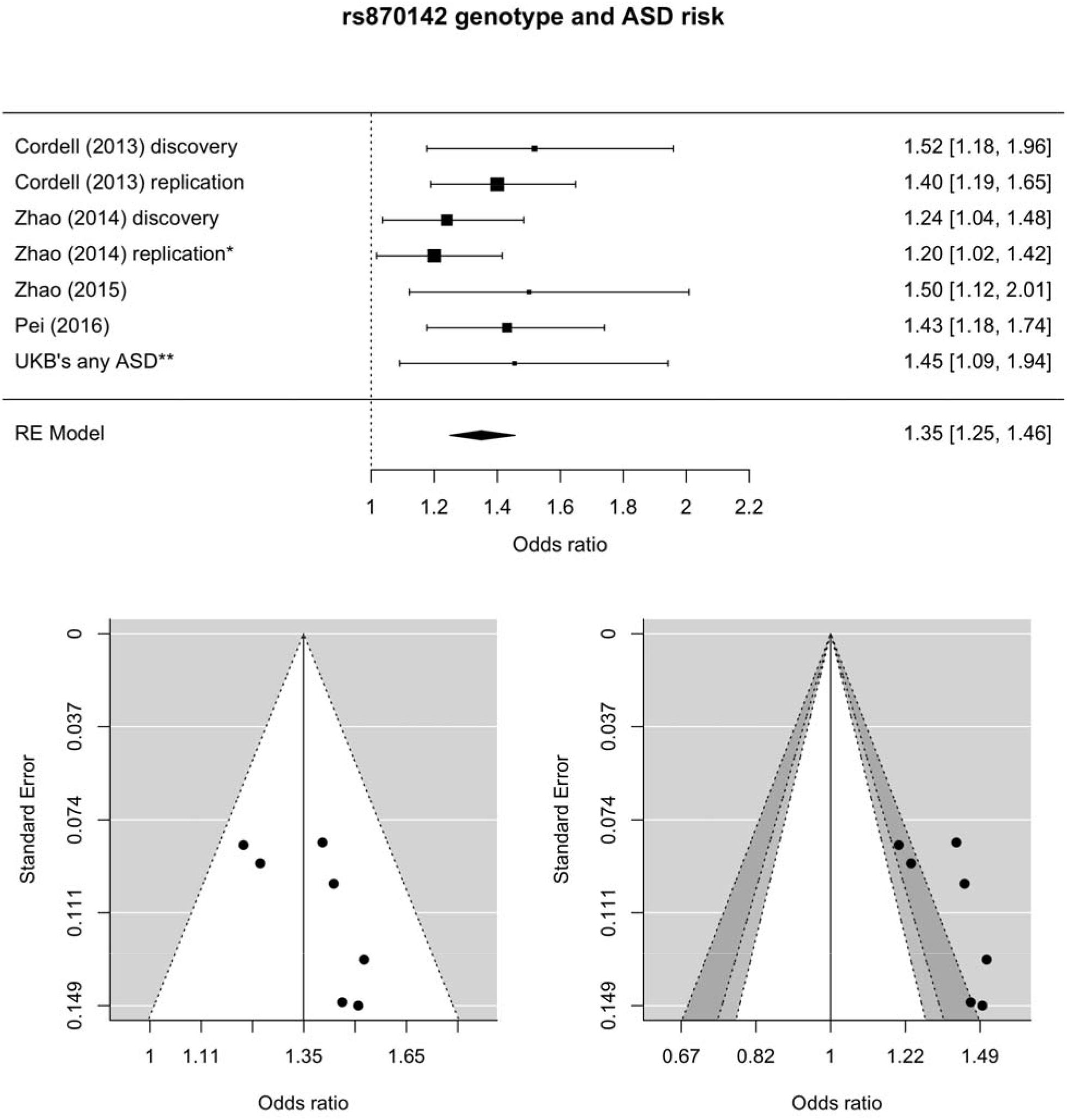
Meta-analysis of rs870142 and ASD risk. Top: forest plot from random effects meta-analysis. Bottom: funnel (left) and contour-enhanced funnel (right) plots. Notes: scale for the observed outcome (x-axis) is odds ratio; *, the replication cohort in the study by Zhao et al. ^7^ had genotypes for rs16835979 instead of rs870142 (they are in strong linkage disequilibrium); **, cases included in this analysis were individuals identified as having ASD with or without another CHD. Light and dark grey shades on the contour-enhanced funnel plot represent 90-95% and 95-99% pseudo-confidence interval regions. Abbreviations: RE, random effects (meta-analysis).

The present findings should be interpreted with close attention to the demographic characteristics of the sample, particularly regarding the age of the participants and the prevalence of CHD in the UK Biobank. With the current diagnostic definitions, there are 821 adult CHD cases (any type, including LVOTO and ASD) and 332,788 controls; this gives a prevalence estimate of 0.25%, which is much smaller than the classically reported ~0.8-1% for CHD in newborns ^1^ and is likely related to survival bias (severe CHD cases would not reach adulthood). Despite this observation, effect sizes observed in the adult ASD survivors from the UK Biobank (OR around 1.3-1.45) were similar to the ones observed in pediatric cohorts (OR around 1.4-1.52, additive model ^6^). In addition, the 4p16-CHD associations (either LVOTO or ASD) were not modulated by age in this sample (Figure 1). Thus, the lower prevalence of CHD in the UK Biobank compared to children samples was apparently not explained by just the 4p16 risk alleles. To some extent, this is also supported by the relatively high MAF of the risk alleles (>23%): the large majority of the allele carriers will not exhibit any CHD phenotype. One could hypothesize that this locus modulates aspects of the disease phenotype that do not have severe clinical implications, or that additional factors are needed to cause deep disruptions (e.g., gene-environment or gene-gene interactions). Along the same lines, another hypothesis is that those risk alleles lead to, for instance, small ASDs that close on their own. Genetic protection against other disease phenotypes may also underlie the findings.

To the knowledge of authors, this is the first report on the association between the 4p16 locus and the LVOTO phenotype. While some previous studies on the association between the 4p16 locus and ASD also tested for associations with other CHD types (e.g., study by Cordell et al. ^6^), most of them did not include results on LVOTO patient data. A similar diagnostic patient set with left heart malformations was available in Cordell et al.’s ^6^ data, comprising 412 subjects with coarctation of the aorta, hypoplastic left heart syndrome and aortic and mitral valve anomalies, but there was no significant association with the 4p16 risk alleles. This discrepancy between those results and the current analysis in the UK Biobank might be due to clinical and demographic differences: the former patient group were children sampled at a clinical setting, with potentially severe conditions, whereas the UK Biobank includes only adult survivors from the general population. Also, although statistically significant at *p*<0.05, the strength of the 4p16-LVOTO association had a smaller effect size and higher *p*-value than the statistics observed for ASD in the UK Biobank, potentially indicative of weaker effects than those observed for ASD. Having noted those potential limitations, Ellesøe et al. ^11^ recently showed that distinct groups of cardiac malformations co-occur in families with a history of CHD, and concluded that there might be shared susceptibility genes for different CHD subtypes. In line with that, the present results provide preliminary evidence of shared genetic risk for LVOTO and ASD.

Meta-analysis of ASD from six previously published data and the UK Biobank outcomes had an overall OR of 1.35 for the rs870142 risk alleles in an additive model (95% confidence interval: 1.25-1.46). Although a simple funnel plot and regression test do not suggest evidence of publication bias (Figure 2, left), lack of data points with trend- and non-significant associations on a contour-enhanced funnel plot (Figure 2, right) constitute an indication of biases, which may include diverse factors such as exaggeration of effects due to lack of statistical power ^12,13^. This could have affected a few of the included results. For instance, the power to detect the effect size reported by Zhao et al. ^7^ was not high, with estimates around 60% in their discovery sample (OR of 1.24, given MAF of 0.333 and disease prevalence of 1%), and 53% in their replication cohort (at OR of 1.2). Power was higher in the study by Zhao et al. ^8^ (~69%); however, they reported an unusual MAF of 28.7% in their controls as opposed to the 33% values reported by the other Chinese groups, which may indicate strong differences in ancestry of the participants. The design by Pei et al. ^9^ was well powered to detect the effect they found (OR of 1.431, power close to 90%). Overall, pooled effect size estimates from the meta-analysis should be interpreted with caution, considering the evidence of bias from the contour-enhanced funnel plot. Studies with larger sample sizes are needed to provide better estimates of the underlying effect size.

Some limitations deserve mention. First, since the UK Biobank cohort comprises volunteers from the general population, the disease prevalence might not be representative of the actual rates observed in the clinics (e.g., severe cases might have not participated). As the number of adult CHD survivors is increasing with the advent of new treatments, the prevalences of different CHD phenotypes will likely change in future studies, limiting between-study comparisons. Moreover, since this is a registry-based dataset and the diagnostic pipeline relied on ICD and OPCS codes, the likelihood of misclassifications and the extent of disease heterogeneity cannot be accurately estimated. Datasets from adult CHD patients recruited and phenotyped directly at clinical settings could help confirm and expand the current findings. Regarding the ASD meta-analysis, attention should be paid to the moderate number of available cohorts (seven independent samples), their modest numbers of ASD cases, within-study power and potential between-study diagnostic differences.

Overall, results of this study support an association between the 4p16 locus and CHD, both in pediatric ASD patients and in adult survivors of ASD and LVOTO. Although further research with large and well characterized patient samples is encouraged, only mild effects (OR~1.35) could be expected based on the current evidence. Hence, the clinicopathological significance of these risk alleles does not seem high, and they may only trigger a disease phenotype in the presence of other risk factors. Additional studies are also needed to clarify why the risk alleles are equally frequent in pediatric and adult CHD cases, and to explain why they predispose only to certain CHD subtypes.

## AUTHOR CONTRIBUTIONS

A.C.-P. and J.R.P. contributed to study design, analysis and interpretation of data, and drafting and critical revision of the manuscript. Final approval of this manuscript version: A.C.-P. and J.R.P.

## DATA AVAILABILITY

UK Biobank data can be accessed following guidelines at https://www.ukbiobank.ac.uk/. Analysis scripts and pipelines are available upon reasonable request to the authors.

## ACKNOWLEDGEMENTS

NIH K99HL130523 to JRP.

## REFERENCES

1 Gilboa, S. M., Salemi, J. L., Nembhard, W. N., Fixler, D. E. & Correa, A. Mortality resulting from congenital heart disease among children and adults in the United States, 1999 to 2006. Circulation 122, 2254–2263, doi:10.1161/CIRCULATIONAHA.110.947002 (2010).

2 van der Bom, T. et al. The changing epidemiology of congenital heart disease. Nat Rev Cardiol 8, 50–60, doi:10.1038/nrcardio.2010.166 (2011).

3 van der Bom, T. et al. Contemporary survival of adults with congenital heart disease. Heart 101, 1989–1995, doi:10.1136/heartjnl-2015-308144 (2015).

4 Zaidi, S. & Brueckner, M. Genetics and Genomics of Congenital Heart Disease. Circ Res 120, 923–940, doi:10.1161/CIRCRESAHA.116.309140 (2017).

5 Simmons, M. A. & Brueckner, M. The genetics of congenital heart disease… understanding and improving long-term outcomes in congenital heart disease: a review for the general cardiologist and primary care physician. Curr Opin Pediatr 29, 520–528, doi:10.1097/MOP.0000000000000538 (2017).

6 Cordell, H. J. et al. Genome-wide association study of multiple congenital heart disease phenotypes identifies a susceptibility locus for atrial septal defect at chromosome 4p16. Nat Genet 45, 822–824, doi:10.1038/ng.2637 (2013).

7 Zhao, B. et al. Replication of the 4p16 susceptibility locus in congenital heart disease in Han Chinese populations. PLoS One 9, e107411, doi:10.1371/journal.pone.0107411 (2014).

8 Zhao, L. et al. Association between the European GWAS-identified susceptibility locus at chromosome 4p16 and the risk of atrial septal defect: a case-control study in Southwest China and a meta-analysis. PLoS One 10, e0123959, doi:10.1371/journal.pone.0123959 (2015).

9 Pei, K. et al. Association Between the 4p16 Susceptibility Locus and the Risk of Atrial Septal Defect in Population from Southeast China. Pediatr Cardiol 37, 120–124, doi:10.1007/s00246-015-1248-8 (2016).

10 Skol, A. D., Scott, L. J., Abecasis, G. R. & Boehnke, M. Joint analysis is more efficient than replication-based analysis for two-stage genome-wide association studies. Nat Genet 38, 209–213, doi:10.1038/ng1706 (2006).

11 Ellesøe, S. G. et al. Familial co-occurrence of congenital heart defects follows distinct patterns. Eur Heart J 39, 1015–1022, doi:10.1093/eurheartj/ehx314 (2018).

12 Button, K. S. et al. Power failure: why small sample size undermines the reliability of neuroscience. Nat Rev Neurosci 14, 365–376, doi:10.1038/nrn3475 (2013).

13 Szucs, D. & Ioannidis, J. P. Empirical assessment of published effect sizes and power in the recent cognitive neuroscience and psychology literature. PLoS Biol 15, e2000797, doi:10.1371/journal.pbio.2000797 (2017).

