## Supplementary Information for "Association between the 4p16 genomic locus and different types of congenital heart disease: results from adult survivors in the UK Biobank"

The next material refers to the manuscript:

Authored by:

*Aldo Córdova-Palomera^1^, James R. Priest^1,2,*^*

*^1^Department of Pediatrics, Division of Pediatric Cardiology Stanford University School of Medicine, Stanford, 94304, CA*

*^2^Cardiovascular Medicine, Stanford University School of Medicine, Stanford, 94304, CA*

Page

Supplementary Methods.	2

Supplementary Figure S1.	5

Supplementary Table S1.	6

Supplementary Table S2.	7

Supplementary Figure S2.	8

References (Supplementary).	9

**Supplementary Methods**

*Ethics statement*

De-identified data from the UK Biobank was used. Ethical approval for the UK Biobank was granted by the North West Multi-Centre Research Ethics Committee and the National Health Service (NHS) National Research Ethics Service (ref: 11/NW/0382). All participants provided written informed consent to participate in the UK Biobank study, and all experiments were performed in agreement with relevant regulations and guidelines. Additional information, including details about the study protocol, is available online (<https://biobank.ctsu.ox.ac.uk/>).

*Participants and phenotype definitions in the UK Biobank*

Genetic and phenotypic information for this study was extracted from the UK Biobank cohort (Bycroft, et al., 2017; Sudlow, et al., 2015), a detailed prospective study of >500,000 individuals aged 40-69 at enrollment. Recruitment was conducted between 2006 and 2010; the study collected extensive phenotypic information about the participants, including medical histories ascertained by healthcare professionals through verbal interview, electronic health records (EHRs) with inpatient diagnosis codes from the International Classification of Disease 9 and 10 (ICD-9 and 10) and Office of Population and Censuses Surveys (OPCS-4) diagnosis codes. In this study, CHD definitions and exclusion criteria were elaborated by leveraging information from medical histories, ICD-9, ICD-10 and OPCS-4 codes, in addition to information on age at diagnosis and age at surgery. The phenotype classification procedure consisted of four main steps:

1) identification of all possible cases based on diagnostic and procedural codes;
2) exclusion of unconfirmed cases for those who met criteria for non-congenital etiologies of heart disease (e.g., rheumatic);
3) confirmation of case status for those whose age at diagnose suggested high likelihood of congenital etiology, or exclusion otherwise; and
4) exclusion of control group for participants with CHD-related conditions due to lack of confidence to ascertain their status.

The procedure is depicted in Supplementary Figure S1. This taxonomy for specific CHD subtypes allowed distinguishing LVOTO and cardiac shunts. Briefly, the LVOTO category comprised the following congenital diseases: hypoplastic left heart syndrome, congenital aneurysm of the aorta, aortic atresia, coarctation of the aorta, aortic insufficiency, aortic stenosis and subaortic stenosis. As undiagnosed aortic valve defects can manifest with outflow obstruction late in life, patients with aortic valve defect not codified as having a congenital heart defect were included. In the entire UK Biobank, there were 236 LVOTO cases and 444 cardiac shunts. Further exclusion criteria were applied based on genetic information (e.g., non-European ancestry) as described next.

*Genetic data and discovery strategy*

The latest UK Biobank data release contains genotypes of 488,377 participants. Genotyping was obtained through the custom UK Biobank Axiom array for most participants, except for a subset of 49,950 subjects involved in the UK Biobank Lung Exome Variant Evaluation (UK BiLEVE), who were genotyped using an Affymetrix Axiom Array. Further details about sample stratification and genotyping platforms can be found elsewhere (Bycroft, et al., 2017; Wain, et al., 2015).

Family relatives and non-European ancestry participants were excluded based on genetic features provided with the latest UK Biobank data release (Bycroft, et al., 2017), and sample sizes included in downstream tests were 164 LVOTO cases, 223 ASD individuals and 332,788 controls. Four genetic variants on chromosome 4p16 were included: rs870142, rs16835979, rs6824295 and rs4689904. Association tests were performed with PLINK 2’s Firth’s penalized regression fallback (Chang, et al., 2015), and included age, gender, 10 principal components related to ancestry and a three-level covariate accounting for genotyping batch. Models without genomic principal components produced equivalent results. Furthermore, changes in case/control genotype frequency ratio across age –due to mortality/survival bias– were tested via logistic regression models with interaction terms in R.

*Meta-analysis of new and existing results*

Random effects meta-analysis of previous results and the novel UKB data was conducted in R using the *metafor* package (Viechtbauer, 2010). The DerSimonian-Laird approach was implemented to account for residual heterogeneity in random effects models, which are suitable for study sets with non-identical methods and samples. Between-study differences were assessed by means of heterogeneity/variability indicators: *I*^2^ (total heterogeneity/total variability), *H*^2^ (total variability/sample variability), *τ*^2^ (estimated amount of total heterogeneity) and Cochran's Q-test for residual heterogeneity (Cochran, 1954) (a measure of whether variability in effect sizes or outcomes is greater than expected based on sampling variability). Potential publication bias effects were tested via standard funnel plots and regression tests, and other potential sources of bias (e.g., exaggeration of effect in poorer study designs with small sample sizes (Schulz, et al., 1995)) were evaluated using contour-enhanced funnel plots (Egger, et al., 1997; Peters, et al., 2008; Sterne and Egger, 2001). Finally, when appropriate, power calculations were performed using CaTS (Skol, et al., 2006).

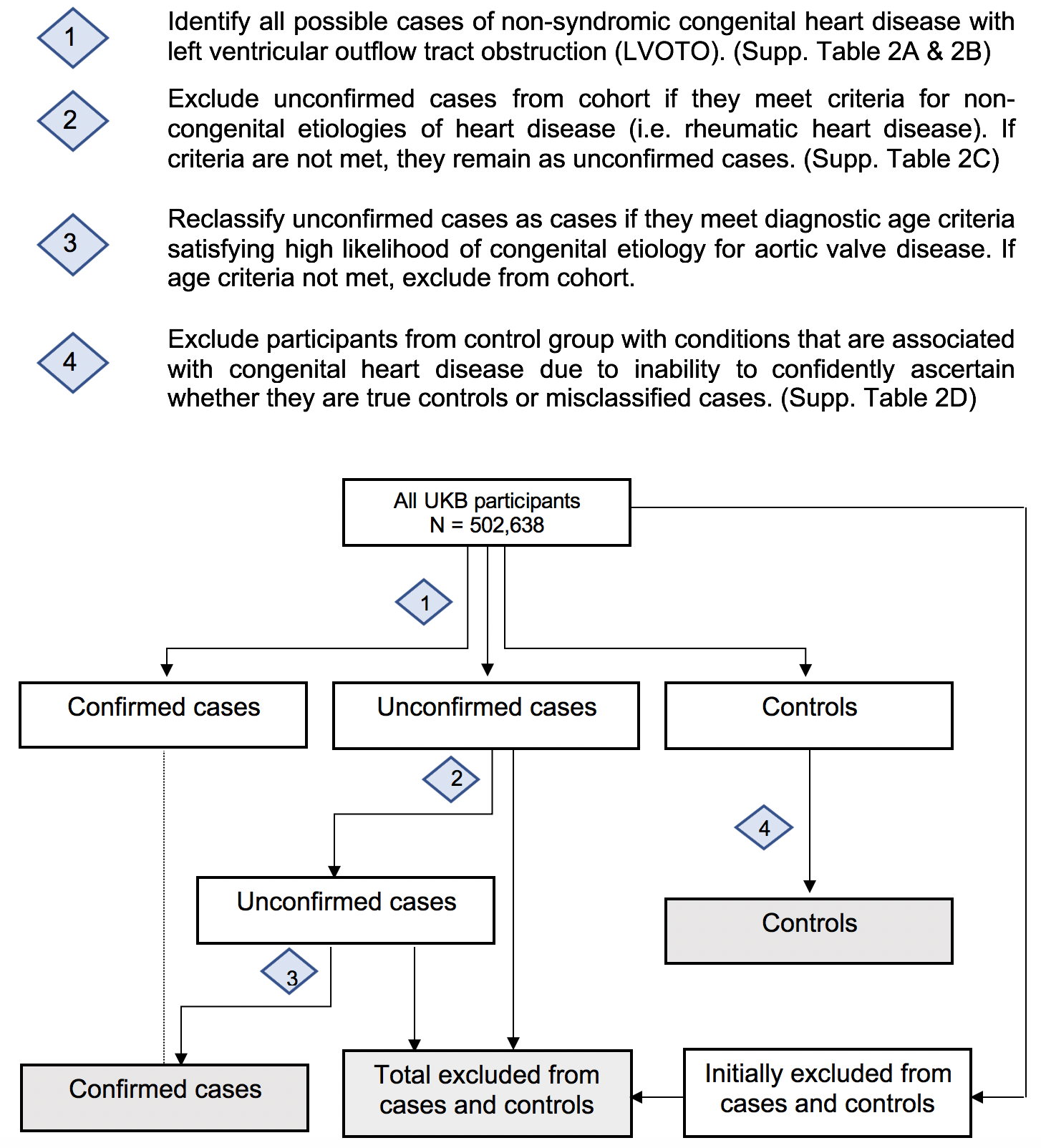

**Supplementary Figure S1.** CHD classification algorithm in the UK Biobank.

|  | **Age (years)** | | | **Gender distribution (*n*)** | |
| --- | --- | --- | --- | --- | --- |
|  | ***Mean (SD)*** | ***Range*** | ***Male*** | | ***Female*** |
| **Controls** | 56.8 (8) | 39-72 | 153,484 | | 179,304 |
| **LVOTO** | 57.6 (7.9) | 40-70 | 111 | | 53 |
| **Any ASD^1^** | 56.9 (8.1) | 40-70 | 129 | | 94 |
| **Pure ASD^1^** | 57 (8.1) | 40-70 | 94 | | 73 |
| **Between-group differences (test statistics)^2^** | | | | | |
|  | *t* | *p* | *X*^2^ | | *p* |
| **Controls vs LVOTO** | -1.2 | 0.245 | 29.8 | | 4.8×10^-8^ * |
| **Controls vs any ASD^1^** | -0.2 | 0.843 | 1.3 | | 0.262 |
| **Controls vs pure ASD^1^** | -0.3 | 0.755 | 0.3 | | 0.585 |
| **LVOTO vs any ASD^1^** | -0.7 | 0.455 | 23.7 | | 1.1×10^-6^ * |
| **LVOTO vs pure ASD^1^** | 0.6 | 0.551 | 18.3 | | 1.9×10^-5^ * |

**Supplementary Table S1.** Demographic information of the participants included in downstream analyses

Notes: ^1^, the “pure ASD” category includes individuals with ASD only, and is a subset of the participants in “any ASD”; ^2^, between-group differences were assessed with either two-tailed *t*-tests (for age) or chi-square (*Χ*^2^, contingency tables with 1 degree of freedom, for sex distributions). Abbreviations: SD, standard deviation; *, statistically-significant p-value.

| **Chr.** | **Position** | **rsID** | **A1** | **A2** | **MAF** | ***n* A1/A1** | ***n* A1/A2** | ***n* A2/A2** | **O(HET)** | **E(HET)** | **HWE *p*** |
| --- | --- | --- | --- | --- | --- | --- | --- | --- | --- | --- | --- |
| 4 | 4614280 | rs6824295 | T | C | 0.2382 | 18735 | 119241 | 190951 | 0.3625 | 0.3629 | 0.5043 |
| 4 | 4628054 | rs4689904 | A | G | 0.2388 | 18977 | 120494 | 192270 | 0.3632 | 0.3636 | 0.5829 |
| 4 | 4635276 | rs16835979 | A | C | 0.2389 | 19032 | 120749 | 192637 | 0.3632 | 0.3636 | 0.5447 |
| 4 | 4648047 | rs870142 | T | C | 0.2379 | 18812 | 120172 | 192760 | 0.3622 | 0.3625 | 0.6457 |

**Supplementary Table S2.** Allelic and genotypic distribution of markers

Abbreviations: Chr., chromosome; rsID, SNP identifier; MAF, minor allele frequency; n, number of observed genotypes; O(HET), observed heterozygote frequency in controls; E(HET), expected heterozygote frequency in controls; HWE *p*, Hardy-Weinberg equilibrium exact test p-value in controls.

**Supplementary Figure S2.** Shift function plots for age and rs870142 genotype for ASD and LVOTO cases, grouped using three genotypes

**References (Supplementary)**

Bycroft C, Freeman C, Petkova D, Band G, Elliott LT, Sharp K, Motyer A, Vukcevic D, Delaneau O, O'Connell J. 2017. Genome-wide genetic data on~ 500,000 UK Biobank participants. bioRxiv:166298.

Chang CC, Chow CC, Tellier LC, Vattikuti S, Purcell SM, Lee JJ. 2015. Second-generation PLINK: rising to the challenge of larger and richer datasets. Gigascience 4:7.

Cochran WG. 1954. Some methods for strengthening the common χ 2 tests. Biometrics 10(4):417-451.

Egger M, Davey Smith G, Schneider M, Minder C. 1997. Bias in meta-analysis detected by a simple, graphical test. BMJ 315(7109):629-34.

Peters JL, Sutton AJ, Jones DR, Abrams KR, Rushton L. 2008. Contour-enhanced meta-analysis funnel plots help distinguish publication bias from other causes of asymmetry. J Clin Epidemiol 61(10):991-6.

Schulz KF, Chalmers I, Hayes RJ, Altman DG. 1995. Empirical evidence of bias. Dimensions of methodological quality associated with estimates of treatment effects in controlled trials. JAMA 273(5):408-12.

Skol AD, Scott LJ, Abecasis GR, Boehnke M. 2006. Joint analysis is more efficient than replication-based analysis for two-stage genome-wide association studies. Nat Genet 38(2):209-13.

Sterne JA, Egger M. 2001. Funnel plots for detecting bias in meta-analysis: guidelines on choice of axis. J Clin Epidemiol 54(10):1046-55.

Sudlow C, Gallacher J, Allen N, Beral V, Burton P, Danesh J, Downey P, Elliott P, Green J, Landray M and others. 2015. UK biobank: an open access resource for identifying the causes of a wide range of complex diseases of middle and old age. PLoS Med 12(3):e1001779.

Viechtbauer W. 2010. Conducting meta-analyses in R with the metafor package. J Stat Softw 36(3):1-48.

Wain LV, Shrine N, Miller S, Jackson VE, Ntalla I, Soler Artigas M, Billington CK, Kheirallah AK, Allen R, Cook JP and others. 2015. Novel insights into the genetics of smoking behaviour, lung function, and chronic obstructive pulmonary disease (UK BiLEVE): a genetic association study in UK Biobank. Lancet Respir Med 3(10):769-81.
